# Humpback whales harbor a highly novel and endemic gut microbiome that adapts to periods of fasting during migration

**DOI:** 10.1101/2025.04.03.647021

**Authors:** Ryan P. Bos, Joe Roman, Michael Fishbach, Suzanne E. Yin, Christine M. Gabriele, Peter R. Girguis

**Author notes:** Corresponding Authors: Joe Roman; Peter R. Girguis. **Competing Interest Statement:** The authors declare no competing interests. **Author Contributions:** J. R., C. M. G., M. F., and P. R. G. conceptualized the research; C. M. G., S. E. Y., and M. F. collected samples; R. P. B., and P. R. G. were involved in data generation; R. P. B., J. R., C. M. G., and P. R. G. processed the samples; R. P. B. processed the data; R. P. B. analyzed data; R. P. B. wrote the first draft of the manuscript. All authors edited and approved the final version of the manuscript.

## Abstract

Humpback whales (*Megaptera novaeangliae*) are cosmopolitan in distribution and most populations migrate between foraging and breeding grounds each year^1,2^. As capital breeders, many humpbacks reduce foraging behavior and potentially fast for a few months of the year^3^. Here, we present a comparative genomic analysis of the gut microbiomes from humpbacks on two breeding grounds: Gulf of California, where feeding is common^4-6^, and Hawai‘i, where feeding is extremely rare^7^. The humpback whale gut microbiome shows unexpected taxonomic novelty, high endemism, and coevolutionary signal. These data also suggest an imbalance in the gut community with foraging reduction, supported by the enrichment of several aerobic metabolism genes, potentially indicative of inflammation, and pathogenesis gene sets used to colonize host tissues, modulate the immune system, and lyse tissues. Pathway reconstruction revealed gut communities shared symbiotic functions including the synthesis of essential amino acids, short chain fatty acids (SCFAs), and vitamins, as well as nitrogen salvaging mechanisms and degradation of biogeochemically relevant substrates like cellulose, chitin, and wax esters (including host chitinases). Complementary sequencing and taxonomic annotation revealed that more Clostridiaceae and Oscillospiraceae genera were in foraging whales, whereas fasting samples were enriched in Erysipelotrichaceae genera, associated with gut disease^8,9^. These data reveal striking changes in humpback whale gut microbiomes between their feeding and breeding grounds, raise the possibility that breeding whales are more susceptible to gut dysfunction or disease, and equally important reveal how humpback whales are likely making varying biogeochemical contributions to upper ocean communities over the course of the year.

## Introduction

Humpback whales (*Megaptera novaengliae*) are among the largest mammals on Earth and play an important ecological role in marine systems^10,11^. These migratory whales are cosmopolitan in distribution^1,2^ and highly metabolically active^12^, consuming predominantly small crustaceans and fishes^13,14^, and as “biological pumps” contribute markedly to nutrient flux in the upper ocean^10^. There are numerous subpopulations of humpback whales that forage in eutrophic waters and seasonally migrate to oligotrophic waters to reproduce^2^, each with differing seasonal ecological impacts. Humpback whales are capital breeders, accumulating lipid stores in the summer to compensate for the net caloric deficit imposed by prolonged fasting experienced during migration^3^. During these migrations, feeding is significantly reduced or absent. Humpbacks off the coast of Hawai’i display iconic seasonal fasting^7^, whereas whales in the Gulf of California appear to feed through much of the winter^4-6^. Though recent efforts have explored the effects of body size on the extreme capital breeding strategy of humpback whales we have no knowledge on the implications of fasting to the gut microbiome, to whale gut health, or to their role in biogeochemical cycles. Here, via an unprecedented set of fecal samples from foraging and breeding ground humpback whales, we examine how the gut microbiome, the resident and transient microbial communities in the digestive tract, may supplement host nutrition and respond to reduced nutrient concentrations.

Whales are homeotherms (i.e., endothermic) that possess a multichambered gut^15,16^, which is characteristic of their terrestrial ancestors (artiodactyls)^15^. Because this gut anatomy and physiology is unique among marine organisms^17^, it may host a distinct gut microbiome. Indeed, previous studies have employed amplicon-based sequencing to reveal distinct microbial communities along the cetacean digestive tract with close associations with their hosts driven by differing symbiont membership^18-25^. Another metagenomic study focused on the gut microbiomes of baleen whales^26^, revealing that baleen whales harbor many similarities to their ruminant relatives such as hosting a gut microbiome that appears to have anaerobic fermentative capabilities^26,27^. Metagenomic and *in-vitro* studies of whale digestate suggest that chitin may also be fermented^26,28,29^. Chitin (C_8_H_13_O_5_N) is one of the most abundant biopolymers in the marine environment and comprises ∼10% of the calories in the humpback whale diet^30^. Gut fermentation of chitin among other carbohydrates may be appreciable microbial sources of ammonia (NH_3)_, dihydrogen (H_2_), organic acids, SCFAs, and biological methane (CH_4_), an important greenhouse gas with high radiative forcing that is important for energy metabolism in the ruminant gut^31^.

While the anatomy of the whale digestive tract is well-characterized^15-17^, functional knowledge of the humpback whale gut microbiome is sparse. Gut microbiomes, especially when ample in size and activity^32^, expand the breadth of host metabolism and *vice versa*, mediating the acquisition of nutrients that may otherwise escape digestion. Gut microbiomes are increasingly recognized to play a role in animal and ocean health, play quintessential roles in digestion and absorption^33^, immune system support^34^, nitrogen salvaging^35^, and vitamin synthesis^36^, ultimately maintaining gut homeostasis^37^. Imbalances in gut microbiome composition can result in dysbiosis^38^, metabolic disorders^39^, and reduced immune functioning^40^, which may render the host susceptible to pathogens and disease.

Lipid studies of *Balaena mysticetus* have also revealed that the select gut microbiota are correlated with wax ester abundance in the gut and may be capable of harnessing energy from wax ester degradation^19^. Wax esters are lipids composed of fatty acids esterified with fatty alcohols that sequester 50% of the autotrophic carbon production in the ocean^41^, and they are abundant in the preferred prey of humpback whales^42^. Wax esters can be mobilized as an energy store by microbes^43,44^ and may consequently supplement whale nutrition during periods of fasting in addition to glycogen stores. Indeed, in other baleen whale species, wax esters are digested and assimilated with high efficiency^45,46^, but this remains to be determined for humpback whales. Mammals do not possess wax ester hydrolases^47^, so it is likely that the gut microbiome harbors the necessary enzymes for wax ester degradation.

In the present study, we functionally and taxonomically characterized fecal microbiome samples from foraging and fasting humpback whales on their breeding grounds via amplicon and whole-genome-shotgun sequencing followed by granular annotation. Using extensive gene catalogs, we performed comparative genomic analyses and reconstructed metabolic pathways to investigate the metabolic differences and similarities between foraging and fasting whale gut microbiomes. Ample coverage enabled binning of the first metagenome-assembled genomes (MAGs) from humpback whale gut microbiomes, so we further functionally and taxonomically classified these MAGs to examine their ecological distribution and genomic characteristics. Surprisingly, we find that both fasting and foraging Humpback whale gut microbiomes are highly endemic and remarkably novel. Up to 107 novel lineages found nowhere outside of baleen whales are most closely related to bacterial genomes from terrestrial gut microbiomes that share common ancestry with whales. These taxa harbor both shared and distinct metabolic capabilities likely correlated with organic matter intake and migratory state, with implications for animal health and biogeochemical cycles.

## Results

Whale fecal samples collected in the Gulf of California (n=3) were classified as foraging (North Pacific summer feeding grounds), whereas those collected near Hawai‘i (n=2) were categorized as fasting (winter reproductive assembly site; Figure 1). We recognize that all samples were collected during the breeding season, when energetic demands peak for pregnant and lactating whales and energy loss is likely for most whales^48^, but Gulf of California whales are observed feeding frequently^4-6^, particularly on krill. We therefore use the terms “foraging” and “fasting” as a shorthand but recognize that these terms refer to feeding patterns across a spectrum of behaviors. Whole-genome-shotgun sequencing followed by preprocessing of raw reads and removal of aligned host reads yielded 214,750,414 high-quality reads across all libraries. Additional metagenomic assembly statistics can be viewed in Table S1. Amplicon sequencing followed by preprocessing of raw reads produced an average of ∼30,000 high-quality reads per library. Phylum and family level taxonomic annotations generated by the complementary taxonomic approach of QIIME2 (taxonomy based on 16S hypervariable regions) and Kaiju (taxonomy based on sequence homology) can be viewed in Figure 2. At the phylum level, both amplicon and whole-genome-shotgun sequencing revealed that fasting and foraging whales were dominated by bacterial genera from the phylum Firmicutes (Figure S1). Bulk taxonomy was not statistically different at the family level (PERMANOVA, R^2^=0.4637, *p*=0.1), however, high congruity between annotations showed that fasting samples were enriched by bacterial genera from the family Anaerovoraceae, Bacteroidaceae, Erysipelotrichaceae, Prevotellaceae, and Rikenellaceae whereas foraging samples harbored more Clostridiaceae and Oscillospiraceae (Figure 2).

**Figure 1.**
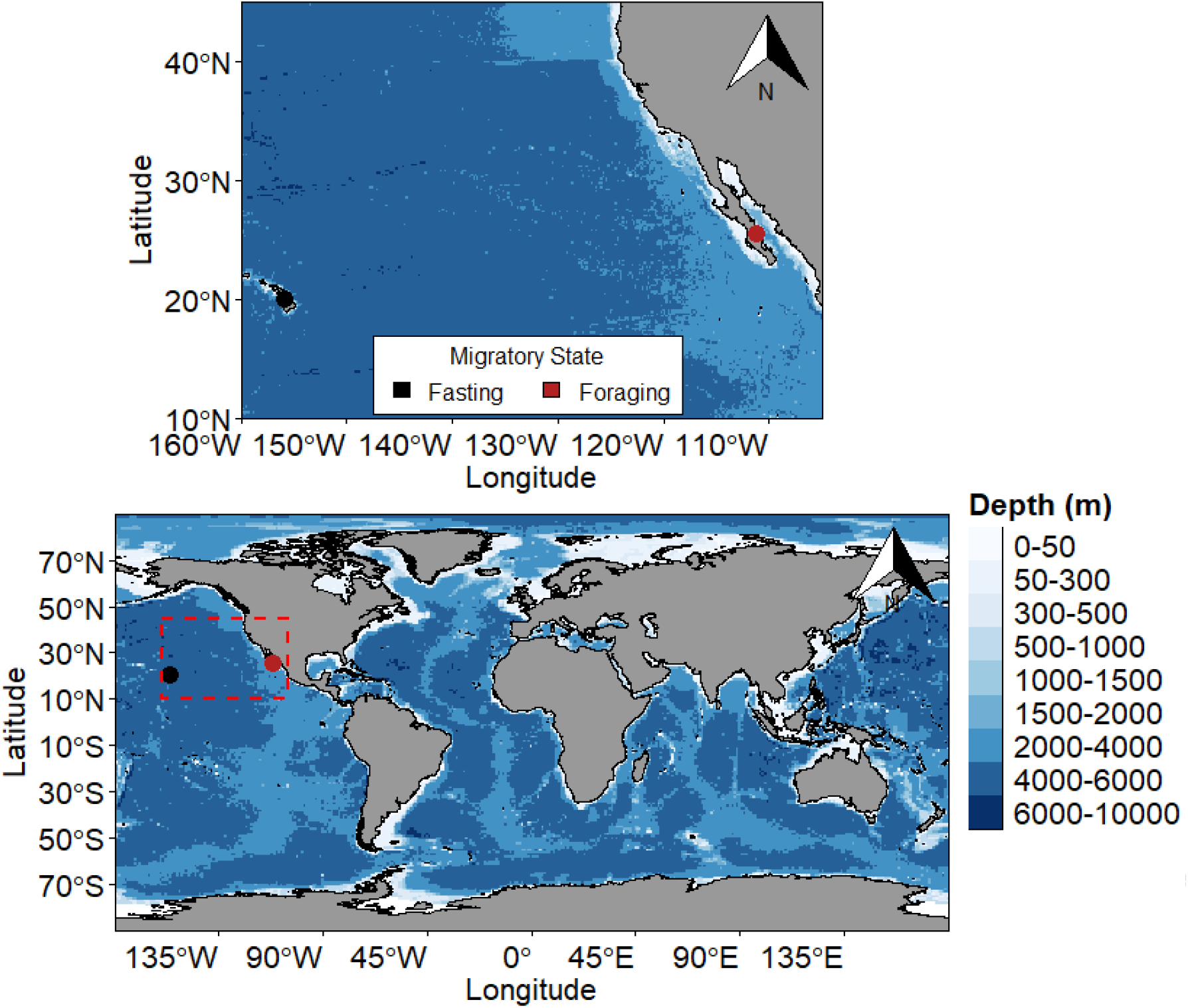
Sampling stations for fasting (black) and foraging (red) whale feces collections. Red dashed rectangle in the bottom panel represents sampling area in the top panel. See Table S1 for more data on collections.

**Figure 2.**
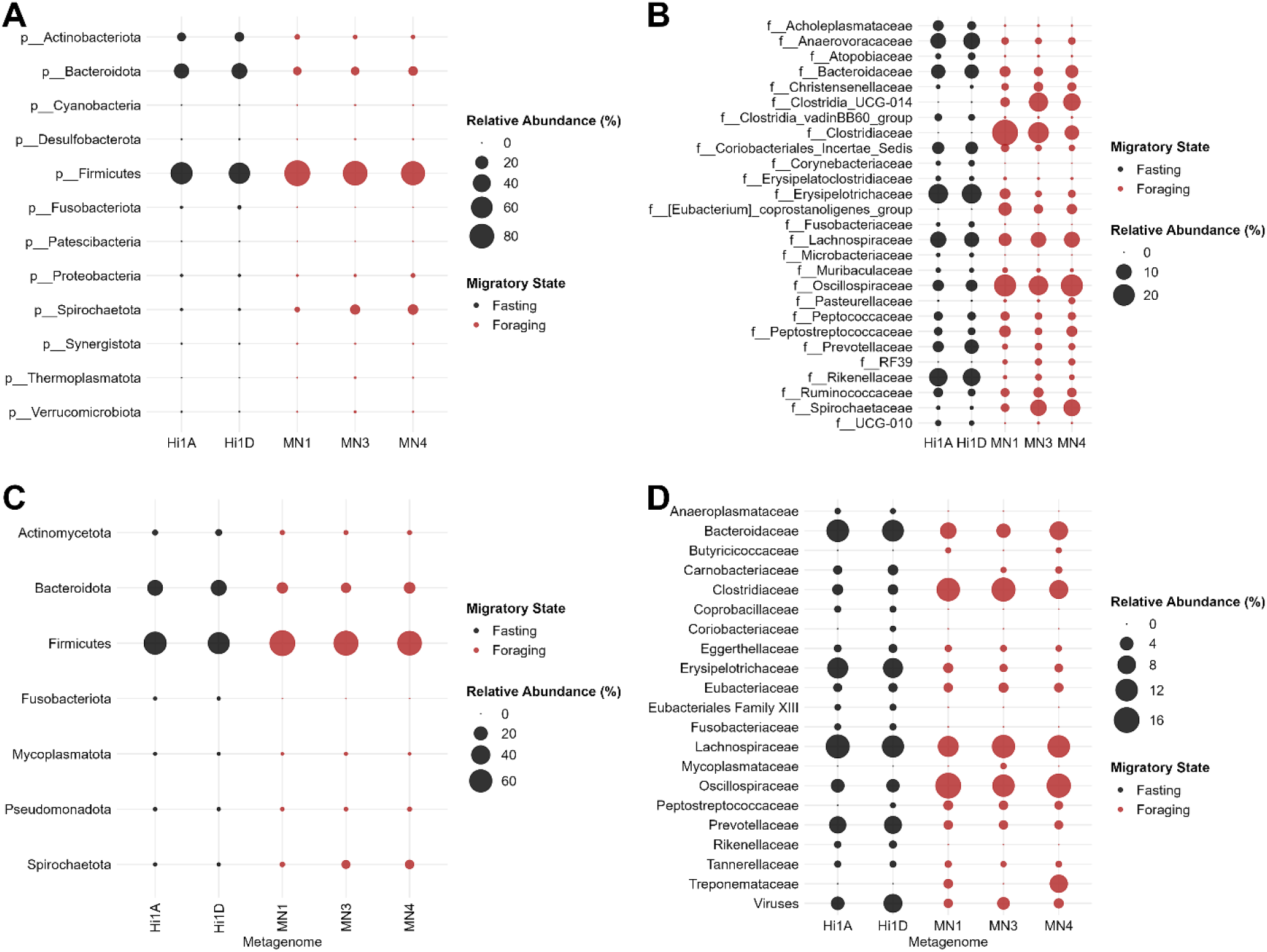
Bubble plots of phylum and family level relative abundances of fasting (black) and foraging (red) humpback whale gut microbiomes for amplicon-based (A, B) and whole-genome-shotgun sequencing (C, D). Relative abundance scales differently between panels.

A total of 80 medium quality and 28 high-quality MAGs were binned and annotated across all samples (Figure 3). Taxonomic annotations and aligned functional annotations with open reading frames, as well as marker gene completion and contamination for MAGs, can be viewed in Dataset 1. Module completion for MAGs is observable in Dataset 2. The normalized median k-mer (k=31) abundance (relative abundance) of our MAGs and the Genbank Bacterial Genomes database (2022, ∼1.2 million genomes) was computed and can be viewed in Figure 3. For example, the relative abundance of MAGs from whale Hi1A accounts for ∼40% of k-mers (k=31), whereas whale MN1 MAGs account for ∼11% of all k-mers (k=31). Exact k-mer (k=31) matching of ∼1.2 million genome signatures from the Genbank Bacterial Genome database represented <0.5% of the relative abundance. When exploring the ecological distribution of our MAGs (≥0.97 cANI), attempted k-mer match containment with millions of metagenomes on SRA suggests that the whale gut microbiome is highly endemic, with 99.1% (n=107) of these novel MAGs being found only in baleen whale gut microbiomes (*Balaenoptera acutorostrata, B. physalus, Eubalaena glacialis, Megaptera novaengliae*) and 0.9% (n=1) found in the Weddell Seal *Leptonychotes weddellii* (Figure 3; Dataset S3).

**Figure 3.**
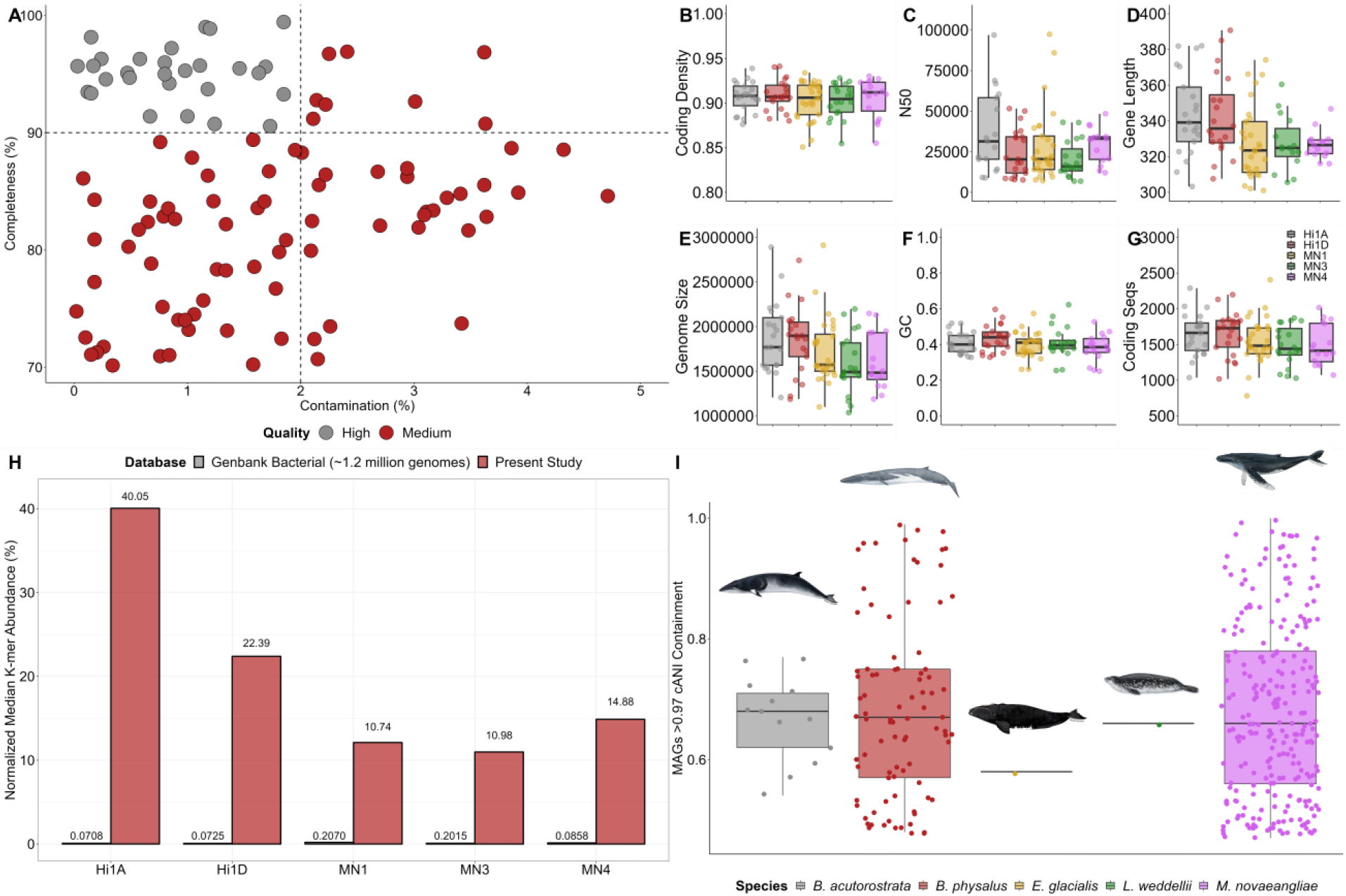
Description and overview, relative abundance, and ecological distribution of metagenome-assembled genomes (MAGs) from fasting and foraging humpback whale gut microbiomes. A) completeness (%) and contamination (%) for high (gray) and medium (red) quality MAGs. B-G) coding density, contig N50, average gene length, genome size (bp), GC content, and number of coding sequences across all whale samples. H) normalized median k-mer abundances (relative abundance) of genomes from Genbank Bacterial database (gray) and MAGs from the present study (red) comprising humpback whale gut microbiomes. I) ecological distribution of MAGs from the present study. See Dataset S1 for MAG annotations and Dataset S3 for containment average nucleotide identity (cANI) calculations. Photo credit: *B. acutorostrata* (NOAA Fisheries); *B. physalus* (NOAA Fisheries); *E. glacialis* (NOAA Fisheries); *L. weddelli* (Voices in the Sea/UCSD); *M. novaeangliae* (Dawn Witherington).

Despite high marker gene completion and low contamination (Figure 3), all 107 MAGs could not be assigned to the species level using GTDB-Tk2. Based on GTDB-Tk2 phylum assignments for our MAGs, *de novo* phylogenetic tree construction based on these phylum level marker genes (Actinomycetota, Bacteroidota, Firmicutes, Spirochaetota, Verrucomicrobiota) and representative RefSeq genomes commonly revealed our MAGs form their own clades (Figure 4). For RefSeq genomes, computations of average nucleotide identity of aligned sequences with our MAGs revealed that all MAGs were <0.89 similar and on average 0.72 ± 0.039 similar to their highest match. For NR, using ribosomal proteins as a guide, MAGs were <0.89 similar and on average 0.76 ± 0.037 similar against the most exhaustive database. Further investigation with NR genomes revealed that the highest ANIb matches were consistently MAGs from the gut environment (n=103), with the top two orders being Artiodactyla (n=37) and Primates (n=37; Figure 4; Dataset S1). Using seven additional hosts and their gut microbiomes, analysis of paired host and gut microbiome phylogenies revealed that gut microbiome composition recapitulated host phylogenetic relationships (Figure S1; Mantel-Test, r=0.791, *p*=0.001, nperm=999).

**Figure 4.**
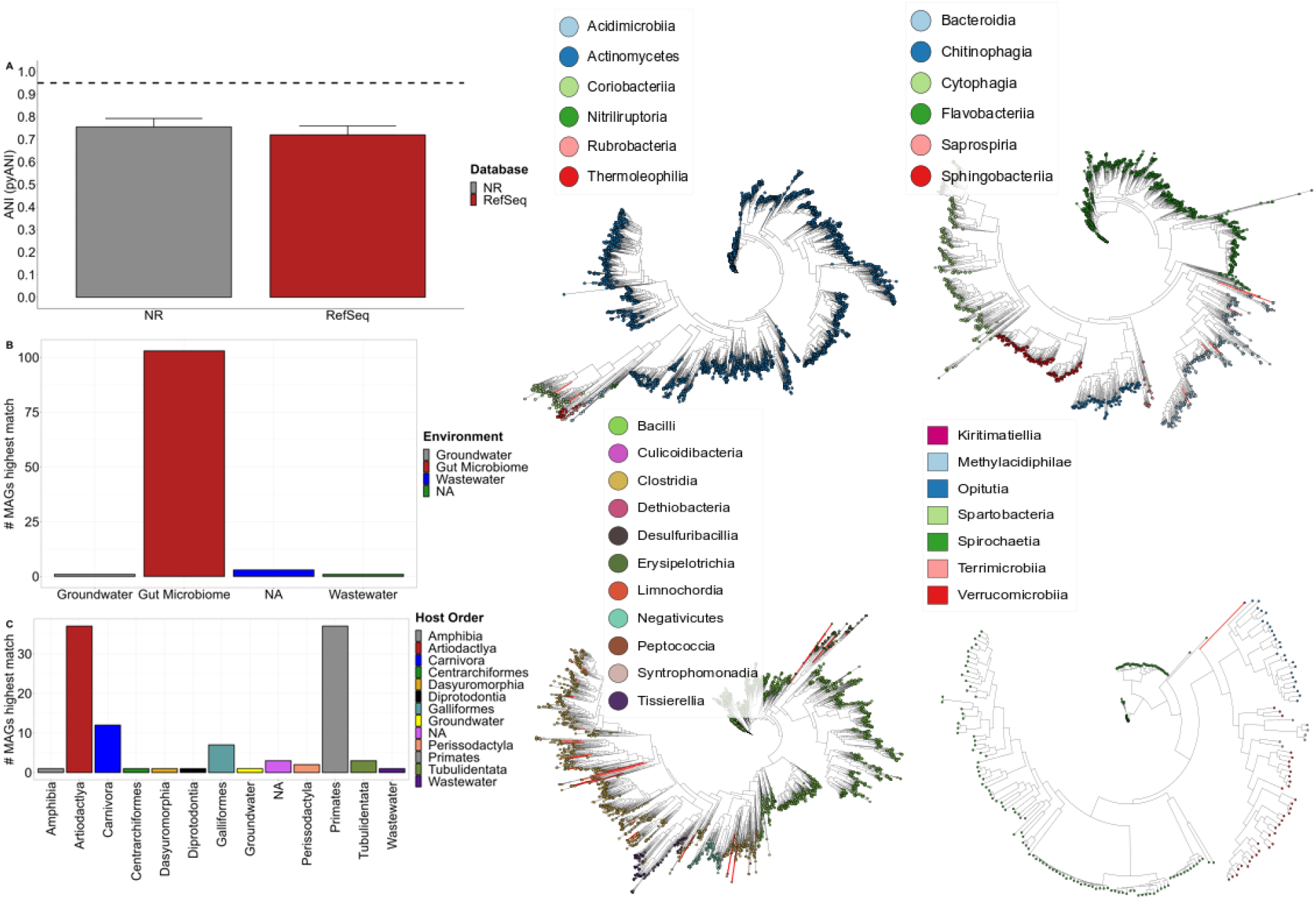
A) Average aligned nucleotide identity (ANI) calculated with pyANI with highest matches from RefSeq (red) and NR (gray). The horizontal dashed line represents the threshold (0.95 ANI match) for taxa to be deemed novel species. B) Count of metagenome-assembled genomes (MAGs) with highest ANI matches across different environments. C) Count of MAGs with highest ANI matches with taxa from gut microbiomes of different host orders and environments. NA depicts host information not available on NCBI. The right panels contain hierarchically clustered phylogenetic trees of novel MAGs from the present study (red branches) and known taxa from RefSeq (black) at the phylum level across different classes. Top left (Actinomycetota); top right (Bacteroidetes); bottom left (Firmicutes); bottom right (Spirochaetota and Verrucomicrobiota). See Dataset S1 for MAG annotations. See Figure S1 for paired host and gut microbiome phylogenies.

Using KOfam annotations, there was no statistical difference between the metabolic potential of fasting and foraging whale gut microbiomes (PERMANOVA, R^2^= 0.9782, *p*=0.1). A total of 20,270 unique gene annotations across four databases (CAZyme, COG, KOfam, and Pfam) were produced by implementing the granular annotation strategy (Figure 5 and Figure S2; Dataset S4-S5). Of these annotations, 12,585 annotations were shared by fasting and foraging whale samples, whereas 7,685 were not (Figure 5 and Figure S2). Pathway reconstruction using KEGG genes revealed 93 nearly complete or completed shared pathways (Dataset S4). Module completion can also be observed in Dataset S2. Essential amino acid and vitamin and cofactor pathways of interest included synthesis of histidine, isoleucine, leucine, lysine, methionine, phenylalanine, threonine, tryptophan, and valine, as well as production of biotin, cobalamin, folate, menaquinone, pantothenate, riboflavin, and thiamine. Other metabolic pathways for amino acid metabolism of interest included arginine and ornithine synthesis, as well as nearly completed urea and ornithine-ammonia cycles. Lipid metabolic pathways were also well-represented across whales including acylglycerol degradation, beta-oxidation, fatty acid biosynthesis, and ketone body biosynthesis. Carbohydrate active enzymes (CAZyme) with chitinolytic potential such as CBM5, CBM37, CBM50, CE4, CE9, GH18, GH19, and GH20 were well represented in all whales. The mucin glycan degradation genes GH29, GH33 GH84, GH85, G89, GH95, GH101, GH129, GH2, GH35, GH42, and GH98 were also well-represented across all whale gut microbiomes (Figure 5; Dataset S5).

**Figure 5.**
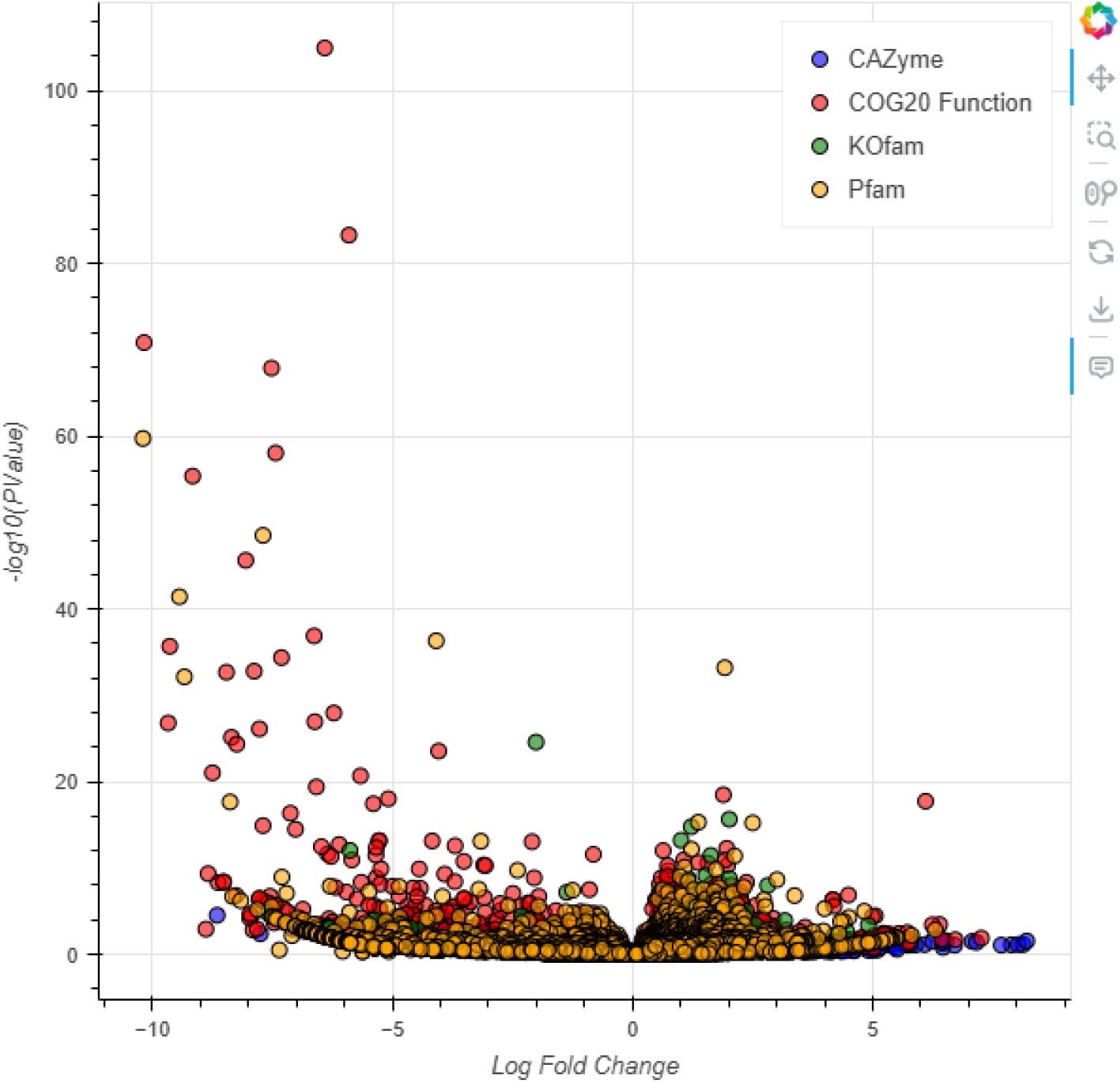
Bokeh plot of a volcano plot displaying differentially present genes (circles) from four annotation databases. Negative log fold changes correspond to enrichment in fasting whale samples, whereas positive correspond with foraging whale samples. Interactive version can be downloaded at https://github.com/Echiostoma/Whale-Gut-Microbiomes.

When comparing differences in metabolic pathways between fasting and foraging whales, foraging whale gut microbiomes harbored either complete or nearly completed pathways for dissimilatory nitrate reduction, methanogenesis with acetate, methanol, and mono-, di-, and trimethylamine as inputs, as well as Reductive citrate cycle and Wood-Ljungdahl pathways (Dataset S4). Foraging whales had significantly higher proportions of genes associated with chemotaxis, motility, sporulation, peptidoglycan synthesis, stress and recombination, and vitamin and cofactor synthesis, whereas fasting whale gut microbiomes comprised enriched gene sets for aerobic metabolism, host cell adhesion and manipulation, iron acquisition, and utilization of diet and host-derived glycans (Figure 5; Dataset S5). Fasting and foraging whales were enriched in 19 and 9 CAZyme families, respectively (Figure S3; Dataset S5).

Polysaccharide degradation potential for all 107 novel MAGs is displayed in Figure 6. Across all MAGs, 18 distinct animal- or plant-based substrates could be potentially used as carbon sources, with chitin, peptidoglycan, mucin, cellulose, starch, milk oligosaccharides, arabinan, and pectin being the top eight most common, respectively. The mean number of carbon sources that could be utilized across MAGs was 6.38 ± 3.24, with a maximum of 13 and minimum of one per MAG.

**Figure 6.**
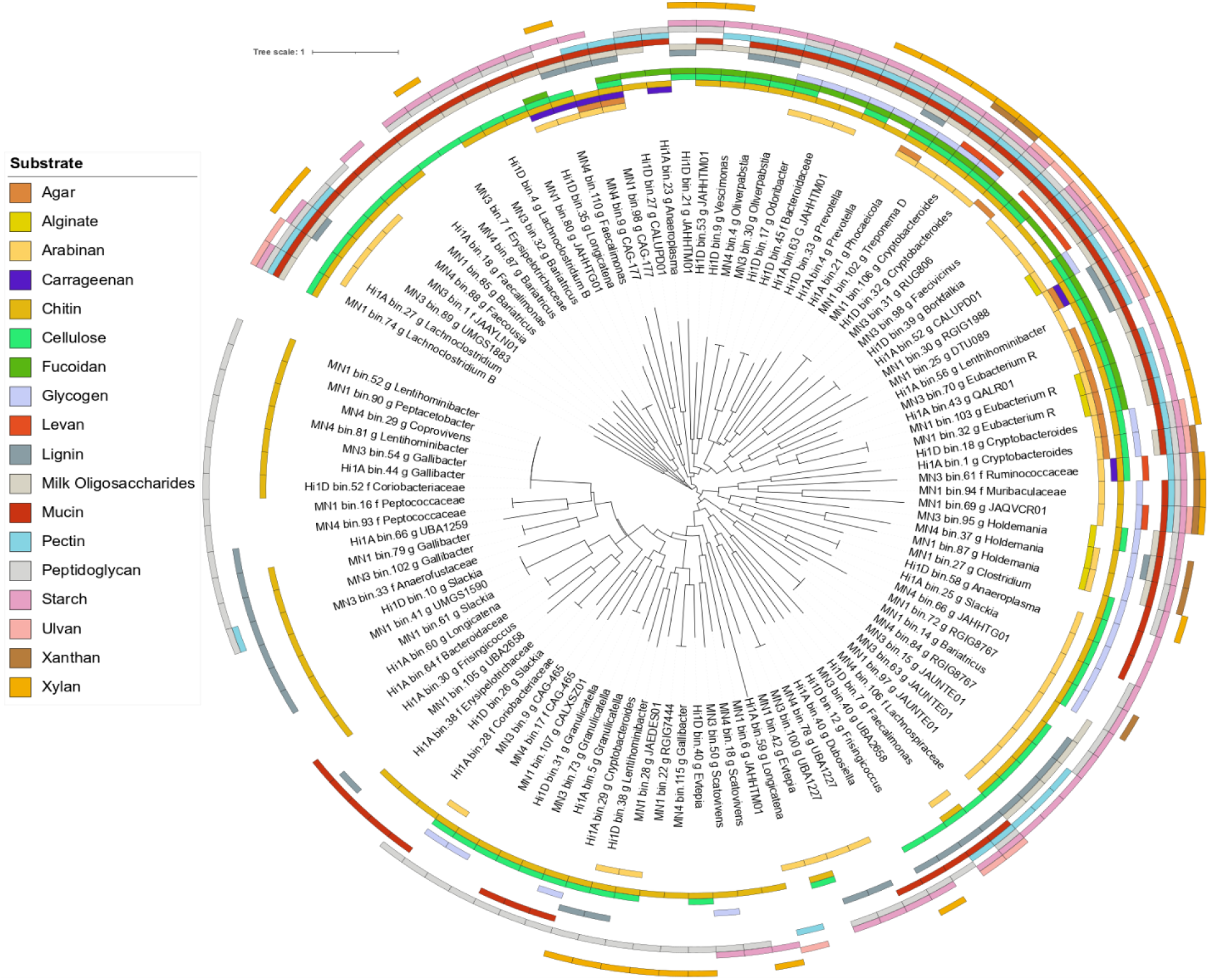
Polysaccharide degradation potential of 18 substrates observable in metagenome-assembled genomes (MAGs) from Hawaiian and Mexican humpback whale gut microbiomes. See Dataset S1 for functional and taxonomic annotations.

## Discussion

Here, we present our analyses of foraging humpback whales and the only known fecal sample to have been recovered from a fasting humpback whale. Humpback whales have been studied for ∼50 years. Necropsies of stranded whales have provided insights into microbial community structure in the gut^19^, but bans on studying or sampling live whales, however, mean there are no data (to our knowledge) that examine changes in gut microbial community structure/function in relation to fasting and feeding. The taxonomic novelty and high endemism of humpback whale gut microbiomes is striking, and similarly, the differences in the taxonomic/functional gut microbiome composition offer new insights into how whale gut microbiomes respond to dietary changes and present new hypotheses about how whales cope with and “recover” from periods of fasting. As humpback whales are endangered, it is important to protect these mammals and their highly endemic communities and functions therein before this microbial biodiversity is lost, potentially before it is even discovered.

### Unexpected Ecological Distributions, Taxonomic Novelty, and Phylosymbiotic Echoes

Genome-resolved metagenomics of fasting and foraging humpback whale gut microbiomes revealed immense taxonomic novelty and restricted ecological distributions of abundant genomes therein (Figures 3-4; Dataset S1). Using annotated ribosomal proteins to inform our searches, comparisons against NR revealed that all 107 MAG lineages fell well below (0.76 ± 0.037) the 0.95 ANI threshold for species identity, suggesting all novel species, and their closest matches were observed predominantly in other gut microbiomes (Figure 3). Moreover, the observation that <0.5% of the normalized relative abundance of genomes could be explained by the Genbank Bacterial Genomes database (∼1.2 million genomes) and species-level matches for 99.1% of our MAGs across >1 million metagenomes on SRA were only observable in baleen whales suggests a highly novel and endemic gut microbiome (Figure 3; Datasets S1 and S3). Although just being discovered now, we posit that the unexpected taxonomic novelty and restricted ecological distribution of members of the humpback whale gut microbiome stems from whales being endothermic homeotherms with terrestrial ancestry, including a multichambered gut and artiodactyl ancestors^15-17^. The former notion is accentuated by evidence of our MAGs having closest ANIb with MAGs predominantly from the guts of taxa from the orders Artiodactyla (close ancestors of whales), as well as Carnivora and Primates (both sharing a distant common ancestor with whales; Figure 4).

Phylosymbiosis is a term used to describe similarities between microbiomes that are driven by host evolutionary history^49^. Phylosymbiotic signal has been previously observed in whales using 16S rRNA data, with Humpback whales having the highest signal^50^. While gut microbiome composition has been shown to be associated with dietary and environmental factors^51,52^, host evolutionary history can also play a profound role in gut microbiome composition^53^. Here, we provide data to support the latter despite clear differences between the diet and environment of the whales and their terrestrial ancestors (Dataset S1; Figure S1). Put a different way, the similarities between gut microbial taxonomic composition recapitulate host evolutionary history and can be used to partly explain our observations (Figure S1). These data may suggest potential phylogenetic inertia, limitations of future evolutionary trajectories imposed by predecessor adaptations. This is not to say that the relationship between host and gut microbiome is suboptimal necessarily, however, as many symbiotic functions are likely performed (discussed below) by the gut microbiome that complement the host and *vice versa*. Alternatively, evolutionary convergence, whereby organisms with distant common ancestors can evolve similar features (gut microbiome functions) independently, may partly explain these trends. Whether coevolution or convergence, both possibilities are likely tied with evolutionary trends in host gut physiology, with potential similarities (e.g., temperature, pH) between mammalian digestive tracts creating similar occupiable functional niches in the gut microbiome^50^.

The restricted ecological distribution of our MAGs could suggest that microbial fitness is decreased with exposure to other host lineages in the marine environment. For example, it could be expected that marine animals might share similar gut microbiomes with other marine organisms like fishes. However, this was observed ∼1% of the time in our dataset (Figure 4), and fish lineages are not closely related to whales. This notion is consistent with natural selection shaping phylosymbiosis, with microbiome transplantation studies demonstrating that key bacterial genera from one host can have significantly reduced fitness in a new host^54^. This may insinuate that the mechanisms of transmission from one generation to the next may be partly explained by defecation (being in close proximity to other whales), as well as calving/lactation, with a strong degree of direct mother to calf microbiome transmission versus colonization from the surrounding environment. The former could potentially lead to more sharing of gut microbiome taxa between baleen whale species for which we predominantly see species-level matches of our MAGs in (Figure 3I).

### Key metabolic differences between fasting and foraging Humpback whale gut microbiomes

The data shown here reveal that a key difference in metabolic potential between actively foraging and fasting was that foraging whale gut microbiomes had more pathways represented for biological methane production, including intermediate pathways of acetoclastic and methylotrophic metabolism (Dataset S4). Methanogenesis has been shown to be correlated with organic matter concentrations^55^, and this observation is consistent with increased organic matter intake (and likely increases in biological methane production) in their feeding grounds. The rates of methanogenesis by this community remain to be determined, though they are plausibly elevated as whales maintain higher internal temperatures and methanogenesis is known to increase with temperature^56^. Moreover, foraging whales had higher representation of Clostridiaceae and Oscillospiraceae (also known as Ruminococcaceae) genera, predominantly obligate anaerobes that play probiotic roles in the gut microbiome (Figure 2). These roles include production of SCFAs like butyrate, which can be important for metabolic cross feeding by microbes and maintaining hypoxic gut conditions^57,58^. Moreover, Clostridiaceae can strengthen the immune system and reinforce the intestinal barrier, providing protection from opportunistic pathogens^59^. Clostridiaceae are also capable of sporulation, which enables these microbes to withstand inauspicious environmental conditions (e.g., lower nutrients, temperature, gut pH) and be transmitted more easily between host organisms^60^. Sporulation gene sets were significantly enriched in foraging whales (Figure 5; Dataset S5). These endospores may enable immune system evasion^61^ and may consequently be relevant to re-establishing quintessential gut functions after periods of migration and fasting. While the hypothesis that when there is insufficient food intake to support vegetative growth, spore-formation increases as spore-formers look to transmit themselves to a new host is logical, isolating and quantifying spores between fasting and foraging whales could provide support or lack thereof for this claim.

Previous work has shown that changes in protein ingestion can lead to an imbalance and therefore shift in microbial community composition of the gut microbiome^62^, and fasting whales have notable differences in taxonomic composition when compared with foraging whales. Namely, Erysipelotrichaceae members (a mix of aerobes and anaerobes) were highly abundant in the fasting whale and have been shown to be highly immunogenic and associated with disease in humans and mice^8,9^, especially after foraging on a lipid-rich diet (characteristic of capital breeders)^63^. Further, there are several genes that are significantly enriched in the Hawai’i whale, suggesting that members of the fasting whale gut microbiome are adapted to colonize host tissues and modulate the host immune system, while potentially being pathogens. The fimbrial surface adhesin *CshA* domains^64^, three immunoglobulin domains^65^, as well as von Willebrand factor binding proteins^66^ suggest host-cell binding capabilities. Von Willebrand factors, in particular, are important for platelet recruitment and coagulation within the endothelium and the binding of bacterial cells to these proteins can enhance inflammation^67^. Ankyrin repeats, used as a form of molecular mimicry by some bacteria, are observable in secreted effector proteins, which can manipulate host cells and enhance virulence^67^, while the MAC/Perforin domains suggest lysis of host tissues and translocation of select molecules^68^. The fasting whale gut microbiome also has significantly higher representation of genes associated with aerobic metabolisms such as cytochrome C oxidases, dioxygenases, heme oxygenases, and microbial globins (Figure 5; Datasets S4-S5). These observations are consistent with the notion that some bacteria reside in the epithelial mucosal layer and consume oxygen, with mechanisms to survive in both aerobic and anaerobic environments.

One of the most striking differences for the fasting humpback whale is the difference in representation of canonical anaerobic metabolisms in foraging whales and aerobic metabolisms in the presumed fasting whale (Figure 5; Dataset S5). In general, mammalian guts have a decreasing intestinal oxygen gradient from the muscle tissue (where it can be quite high) to the lumen (where it is often low). The colonic region of the mammalian gut lumen is typically devoid of oxygen^69^. This gradient is important for gut functioning^70^, in particular retaining obligate anaerobic bacteria. Observations of an enrichment of aerobic metabolism genes in the fasting whale is consistent with one or more of the following scenarios. First, decreases in organic matter consumption by fasting whales, paired with presumed smaller microbial loads, may enable oxygen concentrations to linger in the gut, thereby enabling the coexistence of microaerobic and anaerobic microbes^71^. Second, the Hawaiian whale might be experiencing gut-tissue inflammation and be producing reactive oxygen species^72^. Third, opportunistic pathogenic bacteria may have invaded the endothelium and can “steal” terminal electron acceptors (host oxygen) for aerobic metabolism^73^. The most parsimonious explanation is that decreased organic matter consumption (e.g., fasting) has led to the observed changes. This has been observed in the human gut microbiome, where normal colonocyte metabolism during feeding yields high oxygen consumption to create a hypoxic environment suitable for beneficial obligate anaerobic bacteria^74^. Alteration of colonic epithelium, resultant from starvation, for example, increases epithelial oxygenation, thereby driving an expansion of facultative anaerobic bacteria, a hallmark of gut dysbiosis. Presumably, when returning to foraging consistently after migration, the colonic epithelium will begin to reestablish a hypoxic gut environment and restructuring of the gut microbiome will occur.

### Shared gut microbiome functional attributes among feeding and fasting humpback whales

In other animal gut microbiomes, bacteria produce vitamins^75^, as well as essential amino acids, that animals otherwise need to acquire through diet, that can be absorbed by the host^76,77^. Mucin-derived sugars formed from the breakdown of mucosal glycans represent a consistent source of nutrients for the gut microbiome^78^, which can be used to produce SCFAs^79^, that can control, to some extent, host processes like immune function^80^ and gut barrier function^81^, including reduction of inflammation, while also serving as an energy source. The observation that completed pathways for many essential amino acid and vitamin synthesis and acetate production, in particular, as well as mucin glycan degradation genes were shared across all whale gut microbiomes suggests that the gut microbiome may supplement host nutrition and be especially important during periods of prolonged fasting (e.g., migration).

SCFAs produced by fermentation also play a role in regulating nitrogen supply in gut^82^. The digestive interplay between the host and gut microbiome facilitates nitrogen salvaging, an important evolutionary mechanism of ruminants, whereby toxic ammonia produced from amino acid catabolism in the gut can be transported and broken down in the liver to create urea for excretion^35^. A substantial portion of urea produced by the liver, however, reenters the gut and can be utilized by bacteria to create amino acids, which can be reabsorbed by the host^35,83^. Although counts of microbial ureases, enzymes traditionally involved in nitrogen salvage, were minimal (Dataset S4), the ornithine-ammonia and urea cycles were nearly complete or complete for all whale gut microbiomes (Dataset S4). These observations, paired with the observation that Amino Acid Metabolism constituted ∼7% of COG annotations (Figure S4), suggest that ammonia and urea-N moieties can be created and recycled by the gut microbiome. The ornithine-ammonia cycle has also been shown to be a metabolic adaptation of bacteria to nitrogen fluctuations^80^, which may be important for both gut microbiome and host during periods of prolonged fasting.

Gut microbiome influence spans far beyond the organismal level and extends to the ecosystem level. In terms of carbon cycling, the role the gut microbiome plays in polysaccharide turnover is accentuated by the abundance of our MAGs and their polysaccharide degrading potential (Figure 6). For example, plant polysaccharides like agarose, carrageenan, and fucoidan can compose 50% of the dry mass of macroalgae^84^ and whales do not harbor specific enzymes for degradation of these compounds, yet many of the abundant members of the gut microbiome do possess this molecular machinery. In contrast to plant polysaccharide degradation, metabolites produced from the degradation of chitin may be especially relevant because of their dietary contribution^30^ and because the gut microbiome and host share functional redundancy regarding chitin degradation. Indeed, the host genome possesses acidic mammalian chitinases, which have been shown to degrade chitin in mouse guts^85^, and chitin degrading potential is observable in 86.1% of MAGs, making it the most common polysaccharide that can be potentially degraded (Figure 6). The microbial contribution to chitin degradation in the gut microbiome warrants further investigation as it is among the most abundant marine polysaccharides globally.

The gut microbiome also likely increases the transfer efficiency of energy stored in wax esters. The study by Miller et al.^19^ characterized the lipidome of *Balaena mysticetus* individuals and revealed that wax esters were the most abundant lipids, with a decrease in abundance along the digestive tract. The question that remains is whether the whale, gut microbiome, or both can digest wax esters. Although mammals are generally not capable of degrading wax^47^, the gut microbiome might provide energy stored in wax esters that would otherwise be inaccessible to the host as it is equipped with carboxylesterases, enzymes that hydrolyze ester bonds, as well as many GDSL-like lipases, which can degrade wax esters and are known to affect the deposition and integrity of cuticular waxes in plants^86^. Once ester bonds are hydrolyzed, the gut community is equipped with fatty acid kinases and binding subunits, short-chain and long-chain fatty acid transporters, and the completed beta-oxidation pathway for fatty acid degradation (Dataset S4). Bile acids produced by the host may solubilize intermediate lipid compounds from wax ester degradation^87^ and allow for some form of assimilation. As fatty acid degradation is important for fermentation in gut Firmicutes and can enable synthesis of ketone bodies^88^, alternative energy sources such as wax esters may also be usable as a long-term energy store for bacteria and host during periods of reduced nutrient supply^89^.

## Conclusion

Changes in foraging in humpback whales leads to a marked change in gut microbiome composition. Given the observed changes in community composition and genomic functional potential, fasting whales likely have decreased organic matter supply in the digestive tract and a subsequent change in oxygen concentrations, which leads to a potential increase in inflammation and pathogenesis gene sets. Upon return to foraging, gut hypoxia and gut function are likely restored. The remarkable taxonomic novelty and highly endemic gut microbiome communities was surprising. As mentioned, these gut communities likely play an important role in nutrient transfer between feeding and breeding grounds and global biogeochemical cycles, though this process remains unquantified^48,90^. Studying whale gut microbiomes affords a unique opportunity to address long-standing questions about gut microbiome ecology and evolution, providing an unprecedented opportunity to study how vestigial features such as a chambered gut could be an exaptation, allowing for anaerobic microbial degradation of organics and potentially the evolution of novel endemic lineages that support Earth’s largest animals.

## Methods

Fresh fecal samples were collected from humpback whales off the coast of California and Hawaii using previously described methods^10^ (Figure S2). Fecal samples were stored at -80 °C until ready for processing. Total genomic DNA was extracted from fecal matter using MoBio PowerSoil DNA extraction kit (Qiagen, CA, USA) following manufacturer protocols. Nucleic acids were extracted in triplicates for each sample and pooled for quantification and downstream analyses. Library preparation was done using NexteraXT kit (Illumina, MA, USA) following manufacturer protocols, and DNA was sequenced using whole-genome-shotgun sequencing with a HiSeq instrument at Bauer Core, Harvard University, MA.

The 16S V4 hypervariable region (primers F515/R806) was amplified in triplicates from uniform (1 ng/µl) input template DNA (ultrapure water as a negative control) and Platinum PCR Supermix (ThermoFisher Scientific, MA, USA). Forward and reverse primers were barcoded and appended with Illumina-specific adapters. Thermal cycler conditions were as follows as part of the Human Microbiome Project^91^: initial denaturation at 94 °C (3 m) followed by 30 cycles of denaturation at 94 °C (45 s), primer annealing at 55 °C (45 s), extension at 72 °C (90 s), and final extension at 72 °C for 10 minutes. Sample replicates were pooled and purified using Diffinity RapidTip2 (Diffinity Genomics, PA, USA) pipette tips. Amplicons were sequenced on an Illumina MiSeq using a 500 cycle v2 reagent kit (250 bp) with 15% PhiX genomic library addition to increase read diversity. Negative controls were verified for band absence on a gel but not sequenced.

Demultiplexed 16S reads were processed using Trim Galore! (https://github.com/FelixKrueger/TrimGalore) to remove adapters and low-quality bases. Briefly, in the QIIME2^92^ suite, high-quality reads were merged using vsearch merge-pairs and quality filtered based on q score using default parameters. Using the Deblur workflow, reads were filtered with a 16S positive filter and trimmed to include reads greater than 225 bp. Reads from 16S amplicon-based sequencing were classified using classify-sklearn against the 16S Silva database (v138.1).

Raw reads generated from whole-genome-shotgun sequencing were preprocessed to remove low quality sequences, adapter sequences, and low complexity regions using Trim Galore. FastQC was used to inspect quality of sequences post preprocessing^93^. Before assembling metagenomes, potential host contamination was removed by indexing scaffolds from the *Megaptera novaeangliae* reference genome (GCA_004329385.1) and mapping reads to the scaffolds using bowtie2^94^ (−D 20 -R 3 -N 0 -L 20 -i S,1,0.50). Decontaminated metagenomic reads were taxonomically classified using Kaiju^95^ (taxonomy based on sequence homology) against the nr database (2023-05-10).

Quality controlled reads were assembled using MetaSPAdes^96^ and assemblies were informatically filtered to contain contigs (>300 bp). Assembly statistics such as total length, N50, and GC content were computed using stats.sh in the BBmap suite^97^. Contig databases of metagenomic assemblies were created using Anvi’o v8^98^. Open reading frames were predicted using Prodigal^99^. Prodigal-designated genes were annotated in a granular scheme using CAZymes^100^ (dbCAN2), COGs^101^, KOfam^102^, Pfam^103^ within the Anvi’o platform and subsequently catalogued with scripts available at https://github.com/Echiostoma/Whale-Gut-Microbiomes. KEGG module completion was assessed using anvi-estimate-metabolism, and pathways were reconstructed using the KEGG Reconstruct tool.

Depth of coverage of reads mapping to metagenomic assemblies was calculated using runMetabat.sh and metagenome-assembled genomes (MAGs) were subsequently binned using Metabat2 with a minimum of 2500 bp bins included in assemblies^104^. Marker gene completion and contamination of potential duplicate single-copy genes were assessed using CheckM2^105^, with a medium quality threshold (≥70% completion and ≤5% contamination) and high-quality threshold (≥90% completion and ≤2% contamination) of MAGs for inclusion in downstream analyses. GTDB-Tk2^106^ (v2.4.0, release 220) was used for taxonomic assignments of MAGs. As all MAGs could not be taxonomically assigned at either the genus or species level, class-level marker genes of MAGs and representative genomes from RefSeq were identified, clustered, and aligned to build phylogenetic trees using GToTree^107^ and subsequently visualized in iToL^108^. The average nucleotide identity (ANI) of the alignments between our MAGs and closest RefSeq genome was calculated using pyANI (ANIb), with ANI <0.95 representing novel species. Additional computations of ANI were achieved by querying annotated ribosomal proteins from our MAGs using BLASTp (perc_identity 70; wordsize 18; e-value 1e^-10^) against nr and pulling the genomes associated with the top 5 hits for each gene to analyze similarity with genomes not contained in RefSeq.

Sourmash^109^ (k-mer based taxonomy) was used to evaluate the number of unique k-mers (k=31) from our MAGs that were observable in the Genbank Bacterial Genome database (∼1.2 million genomes), as well as our metagenomes. To determine the ecological distribution of our MAGs, DNA sequences from our MAGs were submitted to the Branchwater Metagenome Query tool to query all public metagenomes on the NCBI Sequence Read Archive (>1 million metagenomes). Matches with a containment average nucleotide identity (cANI) >0.97 were considered to be species-level matches, and normalized relative abundances were calculated with the formulas in eq 1 and 2:

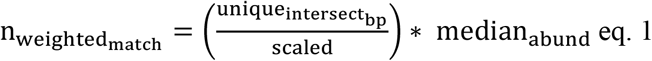

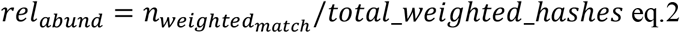

The complete mitochondrial genomes for *Bombus impatiens* (BK063623.1), *Gallus gallus* (AP003580.1), *Homo sapiens* (AB055387.1), *Leptonychotes weddelli* (AM181025.1), *Mirounga leonina* (AM181023.1), *Sus scrofa* (NC_000845.1), *Bos taurus* (V00654.1), and *Megaptera novaeangliae* (AP006467.1) were downloaded from NCBI using the former accessions and aligned using CLUSTALw^110^. Subsequent alignments were imported into r using the seqinr package and converted into a distance matrix and used to create a phylogenetic tree using Neighbor-Joining tree estimation^111^. Whole-genome-shotgun reads of gut microbiomes for all of the former hosts were downloaded from NCBI using fasterq-dump in the SRA Toolkit platform (https://trace.ncbi.nlm.nih.gov/Traces/sra/sra.cgi?view=software). Three replicates of each host gut microbiome and respective host genomes were downloaded and subsequently converted to Bowtie2 indexes as described above. Reads were preprocessed, mapped to their respective host to remove host contamination, and taxonomically annotated as described above. A Bray-Curtis dissimilarity matrix was generated using the total read counts of phylum level annotations of prokaryotes identified in the guts of the former host taxa.

## Statistics

DESeq2^112^, with Benjamini Hochberg Correction of p-values, was used to statistically compare differences in functional metabolic potential between fasting and foraging whales across all four annotation databases. Functional gene catalogs and taxonomic matrices were ordinated by non-metric multidimensional scaling and statistically compared using Permutational Analysis of Variance (PERMANOVA). A Mantel-Test was used to compute the statistical correlation or lack thereof between the host phylogenetic and gut microbiome composition matrices. All statistical calculations were conducted using R^113^. Maps were created using ggOceanMaps^114^ in R.

## Acknowledgements

We acknowledge Neha Sarode for initial processing of samples, including DNA extraction and 16S amplification. We thank Frank Stewart for providing a MiSeq instrument for sequencing. We thank the Harvard University Bauer Core Facility for assistance with library preparation and whole-genome-shotgun sequencing of samples. HMMC sample collected under the authority of National Marine Fisheries Service scientific research permit #15330 issued to Robin Baird, Cascadia Research Collective. Gulf of California whales were collected under the authority of Oficio N° SGPA/DGVS/00282/16 permit.

## Data Availability Statement

Sequencing data generated and analyzed in the present study have been deposited at the NCBI Sequence Read Archive under BioProject ID: PRJNA1239925 and BioSample accession numbers: SAM47492134 – SAM47492138. Code used in bioinformatic workflow can be found at https://github.com/Echiostoma/Whale-Gut-Microbiomes. Supplemental datasets are available at https://doi.org/10.6084/m9.figshare.28719617.v1.

