## Supplemental information for "Humpback whales harbor a highly novel and endemic gut microbiome that adapts to periods of fasting during migration"

**This PDF file includes:**

Figures S1 to S4

Table S1

Datasets S1-S5

### Supplemental Figures and Tables

**Table S1.** Metagenomic assembly statistics for Humpback whale gut microbiome samples.

| Read Library | Number of Reads | Total Length | *N* Contigs | N50 | GC | N Mapped Whale Host Reads | % Whale Host Alignment | Foraging Status |
|---|---|---|---|---|---|---|---|---|
| **Hi1A** | 36906586 | 262176709 | 263389 | 1698 | 0.4193 | 244455 | 0.64 | Fasting |
| **Hi1D** | 48905055 | 225040058 | 215237 | 1911 | 0.421 | 409990 | 0.82 | Fasting |
| **MN1** | 40579628 | 797868294 | 1018833 | 909 | 0.4468 | 43161 | 0.1 | Foraging |
| **MN3** | 49431498 | 813422141 | 1121341 | 788 | 0.4295 | 568453 | 1.14 | Foraging |
| **MN4** | 38927647 | 851870828 | 1145804 | 829 | 0.4371 | 310515 | 0.79 | Foraging |

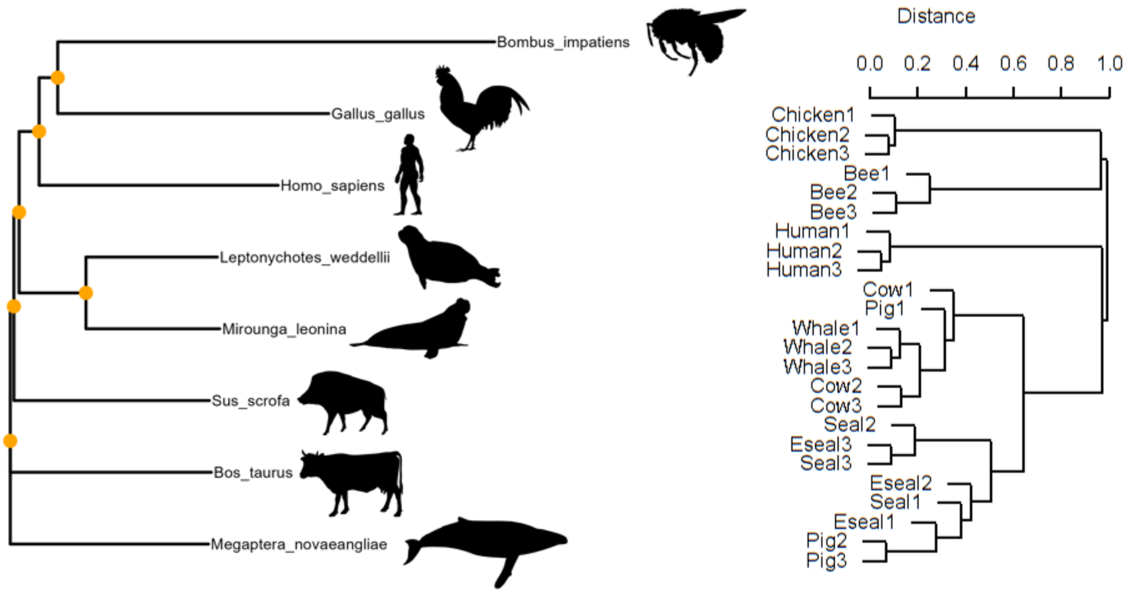


**Figure S1.** Paired host (left) and gut microbiome taxonomic composition (right) of related and distantly related host species. Host phylogeny was created with alignment of complete mitochondrial genomes, whereas gut microbiome phylogeny was constructed with a Bray-Curtis dissimilarity matrix of phylum level taxonomy. Orange circles represent distinct nodes.

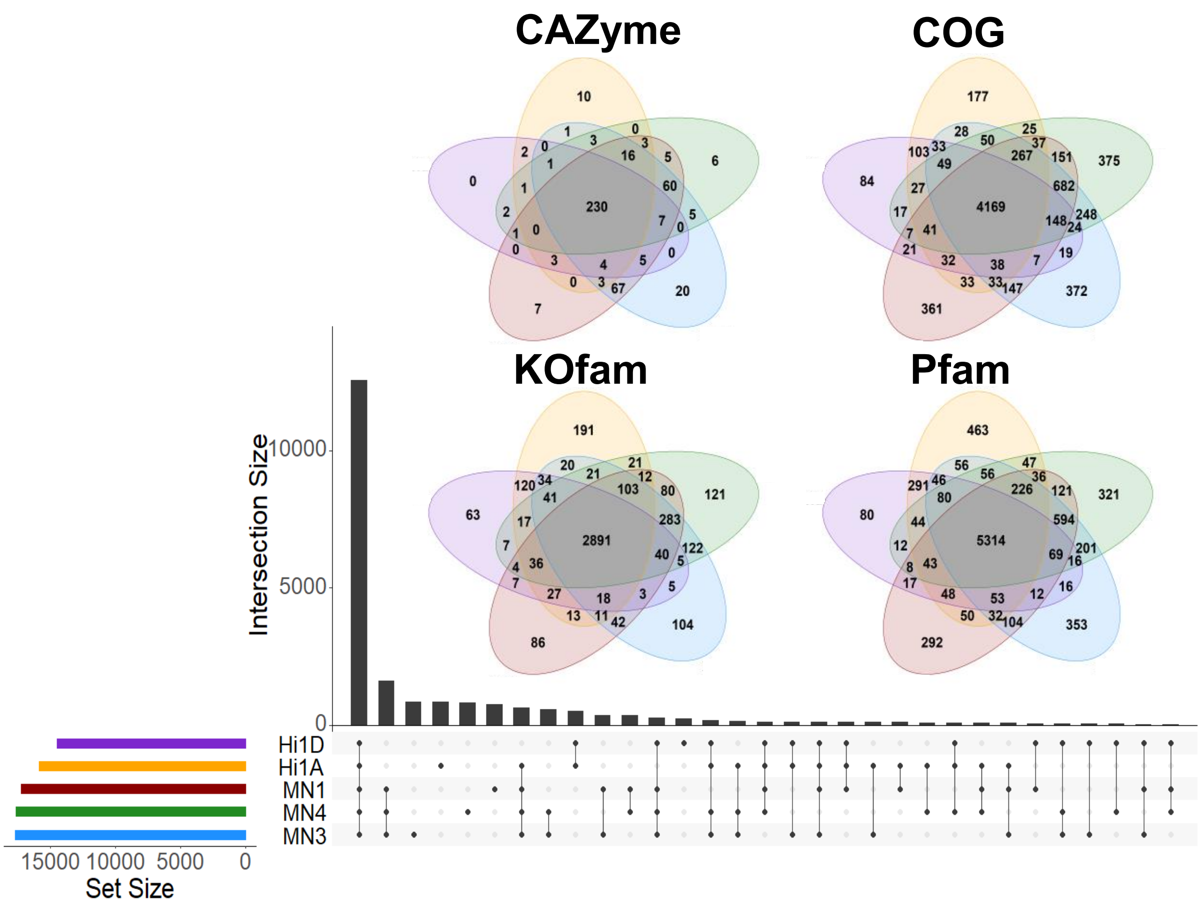


**Figure S2.** Upset plot depicting shared and unique gene annotations across all metagenomes and for each functional annotation database.

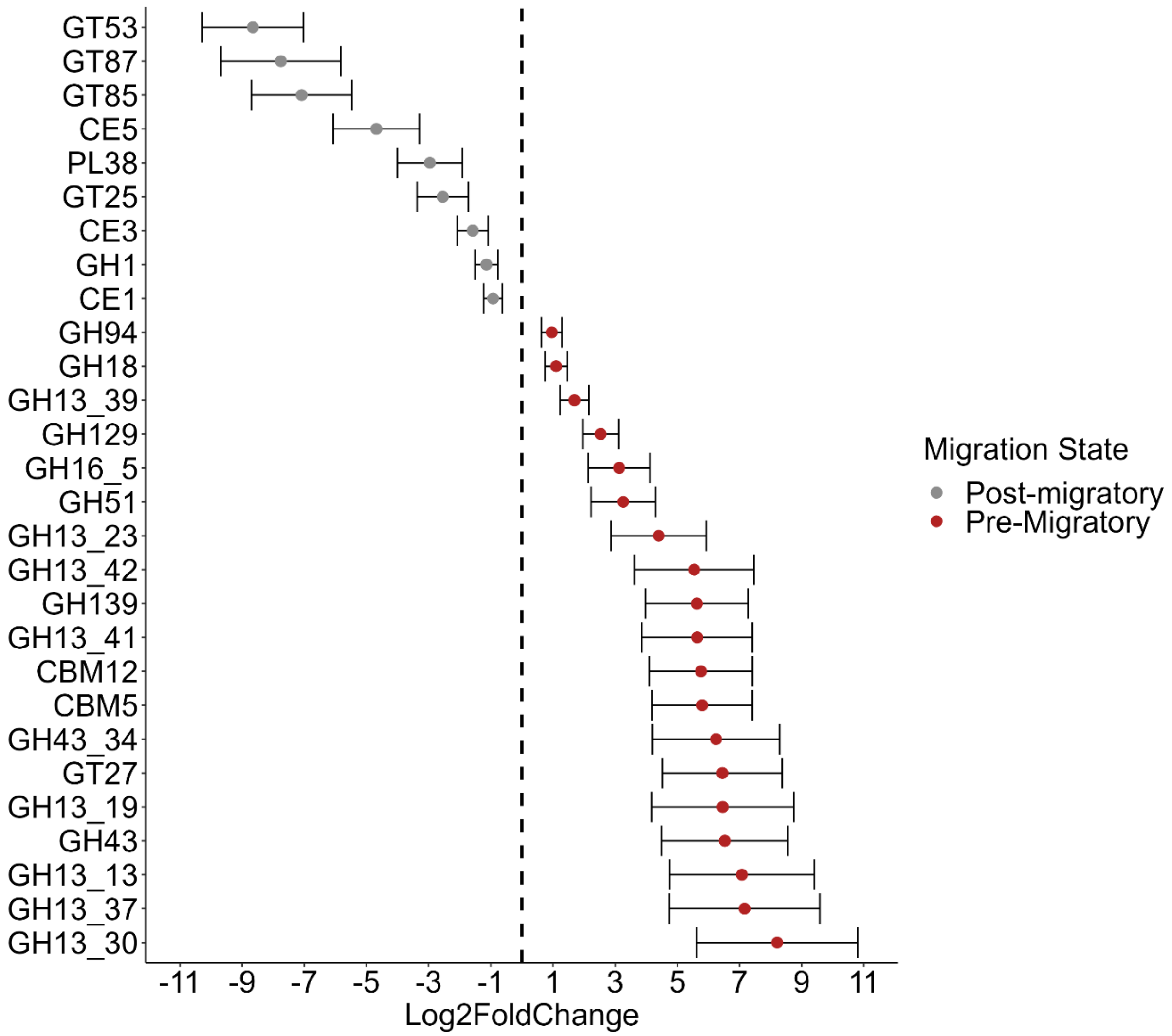


**Figure S3.** The mean $Log_2$ fold changes ± SD of significantly enriched CAZymes between fasting and foraging whales gut microbiomes.

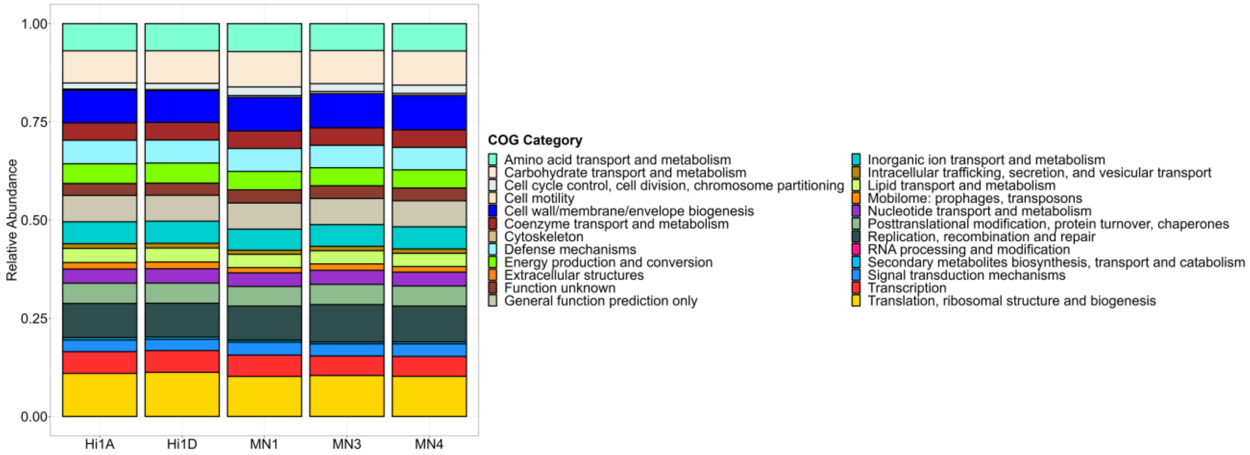


**Figure S4.** Stacked barplot of the relative abundances of Clusters of Orthologous Groups (COGs) categories for whale gut microbiome samples.

#### Additional Datasets

**Dataset S1.** Gene catalog containing functional annotations from CAZyme, COG, KOfam, and Pfam databases.

**Dataset S2.** Pathway module completion for metagenomes and metagenome-assembled genomes.

**Dataset S3.** DESeq2 algorithm outputs for CAZyme, COG, KOfam, and Pfam annotated genes.

**Dataset S4.** Metagenome-assembled genome assembly statistics and aligned open reading frames with functional annotations for all novel lineages.

**Dataset S5.** Ecological distribution of metagenome-assembled genomes.
